# An actomyosin network organizes niche morphology and responds to feedback from recruited stem cells

**DOI:** 10.1101/2023.09.08.556877

**Authors:** Bailey N. Warder, Kara A. Nelson, Justin Sui, Lauren Anllo, Stephen DiNardo

**Affiliations:** Department of Cell and Developmental Biology, Perelman School of Medicine, University of Pennsylvania, Philadelphia, PA 19104, USA; Institute for Regenerative Medicine, University of Pennsylvania, Philadelphia, PA 19104, USA

**Keywords:** Drosophila, Stem cell, testis, niche, feedback, actomyosin contractility, morphogenesis

## Abstract

Stem cells often rely on signals from a niche, which in many tissues adopts a precise morphology. What remains elusive is how niches are formed, and how morphology impacts function. To address this, we leverage the *Drosophila* gonadal niche, which affords genetic tractability and live-imaging. We have previously shown mechanisms dictating niche cell migration to their appropriate position within the gonad, and the resultant consequences on niche function. Here, we show that once positioned, niche cells robustly polarize filamentous actin (F-actin) and Non-muscle Myosin II (MyoII) towards neighboring germ cells. Actomyosin tension along the niche periphery generates a highly reproducible smoothened contour. Without contractility, niches are misshapen and exhibit defects in their ability to regulate germline stem cell behavior. We additionally show that germ cells aid in polarizing MyoII within niche cells, and that extrinsic input is required for niche morphogenesis and function. Our work reveals a feedback mechanism where stem cells shape the niche that guides their behavior.

## Introduction

Stem cells often reside in a specialized niche where they receive signals to balance self-renewal and differentiation. Imbalances in renewal or differentiation can induce tissue degeneration or tumor formation^1,2^, so it is vital to understand mechanisms of niche function. Niches often have a precise morphology^3–6^, but only recently has attention turned to how a niche is shaped and how shape contributes to function^7–9^. Recent work often relied on whole-tissue manipulation or organoid models. Therefore, it is necessary to establish models to interrogate niche morphogenesis with cell-type specificity and in developing tissue.

We have leveraged the genetically tractable *Drosophila* testis, as the niche and associated stem cells are well-defined^10^. This niche is anchored at the testis apex^11^, and is comprised of quiescent somatic cells organized in a reproducible sphere^12–14^. This niche supports two stem cell lineages^10,15^: germline stem cells (GSCs), which produce differentiating germline cells, and Cyst stem cells (CySCs), which produce somatic cells supporting the germline^16,17^.

The niche regulates multiple aspects of stem cell behavior to maintain homeostasis. For example, the niche secretes the chemokine Unpaired (Upd) and activates the Jak-STAT pathway in adjacent cells,^18^ directly promoting CySC renewal and indirectly promoting GSC renewal^19,20^. The niche also regulates GSC division orientation to ensure one daughter remains accessible to renewal signals, while the other is displaced away from the niche and differentiates^21–26^.

While much is known about how the niche supports its stem cells at steady state, little is known about the establishment of a functional niche. Early in embryonic gonadogenesis, the Jak-STAT pathway is active in all primordial germ cells (PGCs) and somatic gonadal precursors (SGPs)^27,28^. Later, the niche forms at the gonad anterior^12,13^, where it selectively contacts and restricts signaling to a subset of cells. Only these cells gain stem-like properties, with PGCs becoming GSCs, and later, SGPs becoming CySCs^27,28^. Because critical niche-stem cell connections become defined concomitantly with niche morphogenesis, it begs the question of how niche shape affects its ability to define and regulate the stem cell pool.

In a major advance, we developed techniques to capture niche and stem cell behavior through live-imaging during this previously elusive period of gonadogenesis^29^. This, together with fixed analyses, has defined two dynamic processes during niche formation: 1) assembly and 2) compaction^12,13^. During assembly, pro-niche cells migrate towards the gonad anterior. Next, during compaction, niche cells reorganize to present a smoothened boundary to the surrounding cells. We have previously shown that errors in assembly lead to defects in niche polarity, quiescence, and function^30^. The requirement for compaction in niche function has not been explored.

Here, we identify a role for polarized actomyosin contractility (AMC) in establishing the final shape and function of the niche. We find that Non-muscle Myosin II (MyoII) within niche cells establishes increased tension along the niche-GSC boundary required to smoothen that interface. When niche structure is altered, we find defects in both signaling and GSC centrosome orientation. Furthermore, compaction timing correlates with the recruitment of actively dividing GSCs, and GSC divisions aid in polarizing niche MyoII required for niche shape and function, likely through a force dependent mechanism. Thus, our analyses identify cytoskeletal mechanics required to form a functional niche and reveal a feedback mechanism in which GSCs shape the niche that guides their behavior.

## Results

### Niche compaction is characterized by changes in niche shape and size

To characterize compaction dynamics, we live-imaged fluorescently labeled gonads at embryonic stage (ES) 16, when niche cells had assembled but not compacted^12,29^. To examine niche contour, we expressed a somatic marker for filamentous actin (F-actin; Six4-Moe::GFP). We previously showed Six4-Moe::GFP permits identification of niche cells due to their increased fluorescence compared to all cells with the exception of, and location opposite to, the male specific SGPs (msSGPs)^12^. To visualize nuclear movements, we utilized a ubiquitously expressed histone marker (His2Av::mRFP1). Imaging onset revealed niche cells at the gonad anterior, forming a jagged contour between themselves and adjacent germ cells (Figure 1A). Within a few hours, niche cells moved closer together and the contour facing the germline adopted a more circular profile (Figure 1A’-A’’’), consistent with previous observations^12,13^.

**Figure 1:**
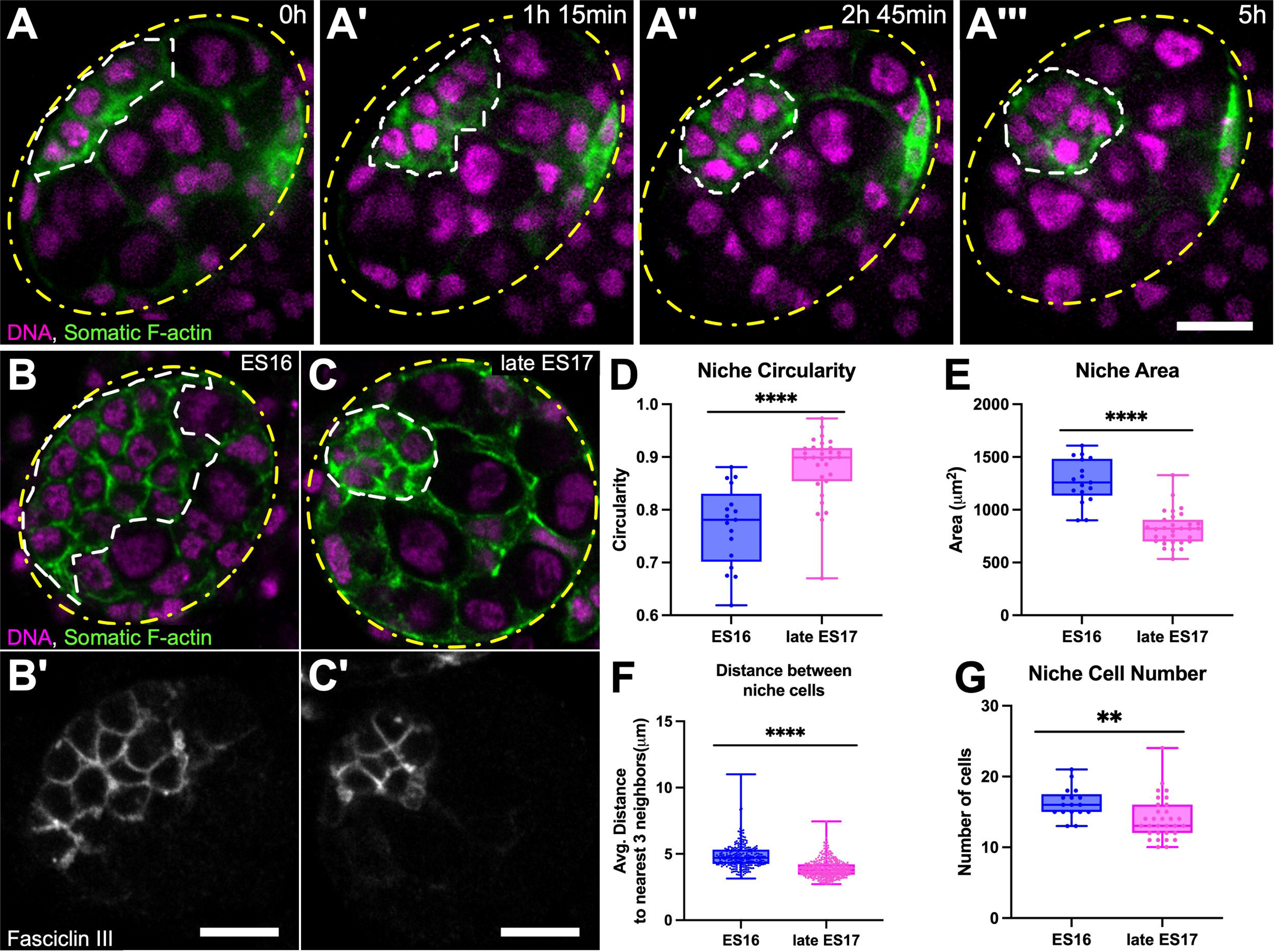
Niche compaction is characterized by a change in niche shape and size A-A’’’) *Ex vivo* gonadal timelapse (5-hour), revealing niche compaction; Six4-Moe::GFP (somatic F-actin, green); His2Av::mRFP1 (nuclei, magenta). A) 5 niche cells in-view reveal an elongated, jagged niche at the anterior. A’) Niche cells rearranged with neighbors, moving closer together. A’’) 6 niche cells in-view present a more rounded contour. A’’’) 7 niche cells in-view are yet closer, presenting a circular contour. B-C) Head-on views of niches from fixed ES16 (B) or late ES17 (C) gonads developed *in vivo;* Six4-Moe::GFP; His2Av::mRFP1. B’-C’) FasIII. D-G) 3D analysis of niches developed *in vivo.* Niche circularity (D) increases post-compaction. Niche surface area (E), niche cell internuclear distance (F), and niche cell number (G) decrease post-compaction (** p <0.01, ****p<0.0001, Mann-Whitney). In D, E, and G, each dot represents one gonad pre- (n=17) and post- (n=31) compaction; in F, each symbol represents one niche cell pre- (n=277) or post- (n=435) compaction. Unless stated otherwise, for all figures: scale bars = 10 microns; images = single Z slice; yellow dashes outline entire gonad; white dashes outline niche.

To confirm our results *in vivo*, we analyzed niche morphology at pre-compaction (ES16) and post-compaction (ES17) stages in fixed preparations using the niche-specific marker Fasciclin III (FasIII) (Figure 1B, B’, C, C‘). We quantified niche contour changes by comparing circularity of the niche outline, where a value of 1 represents a perfect circle and lower values indicate a more jagged contour. Like our *ex vivo* results, niche cells *in vivo* evolved a more circular form from pre- to post-compaction (Figure 1D). 3-Dimensional analysis revealed that niches decrease in surface area (Figure 1E). This was partially explained by niche cells moving closer, as evidenced by a decrease in niche cell internuclear distance (Figure 1F). We also noticed a variable and slight decrease in niche cell number from pre- to post-compaction (Figure 1G). Overall, compaction of the niche occurs both *ex vivo* and *in vivo,* suggesting this process can be used to identify factors required for niche morphogenesis. Below, all live *ex vivo* experiments were performed on ES16 gonads to visualize compaction, whereas *in vivo* fixed experiments were performed on late ES17 gonads for endpoint analyses.

### F-actin and MyoII polarize along the niche periphery during compaction

Due to the role of AMC in many morphogenetic events^7,8,31–35^, we hypothesized that AMC drives niche compaction. To visualize F-actin and MyoII dynamics, we live-imaged gonads containing Six4-Moe::GFP and MyoII regulatory light chain (MyoII RLC) tagged with mCherry (MyoII:mCherry; Figure 2A). We found F-actin localizes along the niche periphery throughout compaction (Figure 2A’). However, MyoII progressively enriches over a few hours during compaction, appearing as puncta along the niche-germline interface (Figure 2A’’, arrow). These enrichment trends were confirmed by quantification along the niche-germline boundary at 0h, 2.5h, and 5h post-dissection, representing early-, mid-, and post-compaction, respectively (Figure 2B-C).

**Figure 2:**
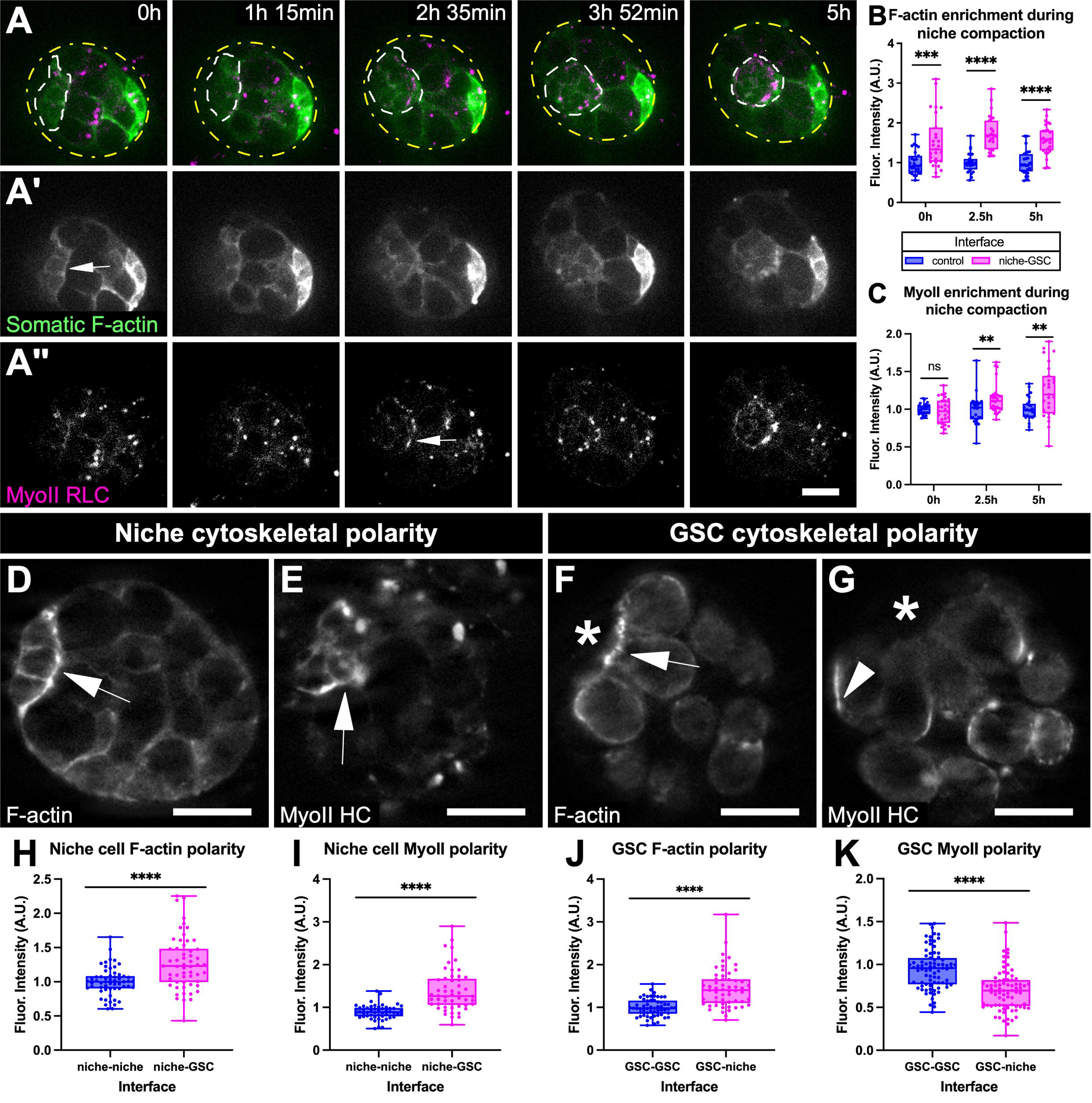
F-actin and MyoII are enriched along the niche-GSC interface during compaction. A) *Ex vivo* gonadal timelapse of niche compaction; Six4-Moe::GFP; MyoII::mCherry. A’) Somatic F-actin. A’’) MyoII. B-C) Quantifications of F-actin (B) and MyoII (C) along niche-GSC interfaces (magenta) relative to non- niche somatic interfaces (blue) at 0h, 2.5h, and 5h post-dissection. Each symbol represents normalized fluorescence along one interface (n=27 for both). D-G) Fixed images of lineage-specific F-actin or MyoII. D) Somatic F-actin (Six4-Gal4 > UAS-F-Tractin::TdTomato.) E) Somatic MyoII (Six4-Gal4::VP16 > MyoII HC::GFP). F) Germline F-actin (Nanos-Moesin::GFP). G) Germline MyoII (Nanos-Gal4::VP16 > MyoII HC::GFP). H-K) Fluorescence along the niche-GSC interface (pink) compared to niche-niche (blue, H-I) or GSC-GSC interfaces (blue, J-K). Equal numbers of niche-GSC interfaces were analyzed compared to respective controls (H: n=57, I: n=52, J: n=57, K: n=79; **p<0.01, ***p<0.001, ****p<0.0001, Mann-Whitney). Arrows and arrowheads = polarization towards or away from niche-GSC interfaces, respectively. Asterisk = niche.

The correlation of MyoII enrichment with morphogenesis suggests a functional role for AMC in niche compaction.

Enrichment along the niche boundary might reflect a symmetric contribution from the niche cortex and the adjacent germline, or an asymmetric contribution from one lineage. To distinguish these possibilities, we independently labeled F-actin or MyoII in a lineage-specific manner (Figure 2D-G). For somatic labeling, we used Six4-Gal4 to drive an F-actin (F-Tractin::tdTomato; Figure 2D) or MyoII heavy chain marker (MyoII HC::GFP; Figure 2E). For germline labeling, we used the germline-specific F-actin marker, Nanos-Moesin::GFP (Figure 2F), and the germline-specific Gal4 driver, Nanos-Gal4::VP16, to express MyoII HC::GFP (Figure 2G). Quantitative analysis revealed F-actin polarizes towards the niche-GSC interface in both niche cells and germline cells (Figure 2D, F, H, J). Conversely, only when labeled within niche cells did MyoII HC polarize to the niche-GSC interface (Figure 2E,G,I,K). The asymmetric recruitment of MyoII in niche cells suggests that tension specifically within the niche cell cortex is required for compaction.

### AMC induces tension along the niche-GSC interface

To test this, we dissected gonads, and identified niches mid-compaction, correlating with when MyoII polarizes along the niche-GSC interfaces (Figure 2C). Using a laser, we severed the actomyosin network either at the niche-GSC interface or at a niche-niche interface (Figure 3A-B). As a proxy for tension, we calculated the initial retraction velocity of vertices flanking a cut by measuring the displacement of each vertex within 5s post-cut (Figure 3D; ref.^36^). This revealed higher tension along niche-GSC interfaces compared to niche-niche interfaces, suggesting a correlation between MyoII polarization and tension. To assess whether the tension along the niche-GSC interfaces is induced by AMC, we used the H-1152 Rho Kinase inhibitor (ROKi) to inhibit AMC, then severed niche-GSC interfaces (Figure 3C-D). ROKi treatment reduced tension along the niche-GSC interface (Figure 3E), confirming that AMC induces polarized tension during compaction.

**Figure 3:**
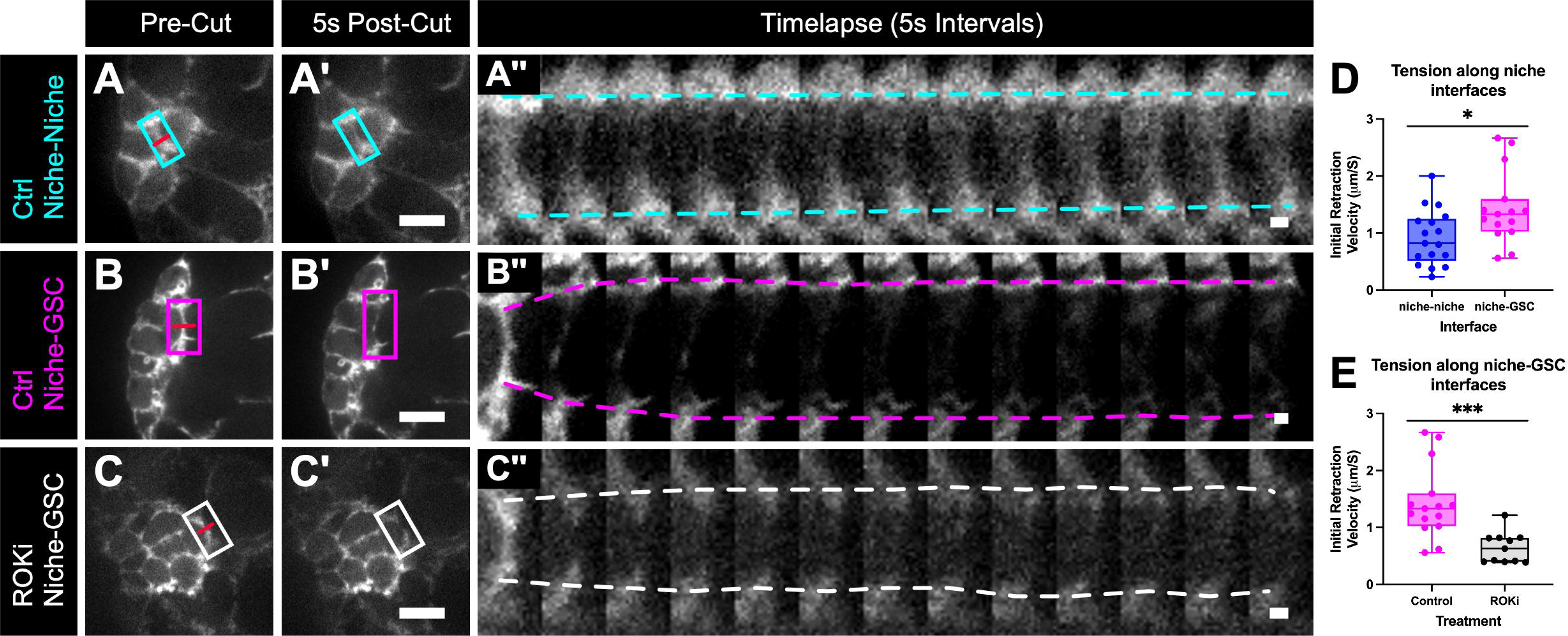
AMC induces tension along the niche-GSC interface. A-C) Six4-Moe::GFP gonads developing *ex vivo.* Outlines = interfaces selected for severing pre-cut (A, B, C; red line = interface targeted) and 5s post-cut (A’, B’, C’). A’’-C’’) Interface pre-cut and montage of 5s intervals post-cut. Dashed lines show displacement of vertices. Scalebars = 1 micron. A) Control niche-niche interface. B) Control niche-GSC interface. C) ROKi-treated niche-GSC interface. D) Initial retraction velocities along niche-niche (blue, n=17) and niche-GSC (magenta, n=15) control interfaces show higher tension along niche-GSC interfaces. E) Initial retraction velocities along niche-GSC interfaces in controls (magenta; same as D) and ROKi- treated gonads show tension is decreased upon AMC inhibition (black, n=11; *p<0.05, *** p<0.001 Mann- Whitney).

### AMC is required for niche morphogenesis

To address the role of polarized AMC in niche morphogenesis, we monitored compaction live in gonads cultured in the presence or absence of ROKi (Figure 4A-B). Untreated niches generally exhibited an increase in circularity over time (Figure 4C; average increase of 20%; median, 8%). Conversely, ROKi treatment administered at compaction onset resulted in mixed circularity changes, with many niches exhibiting decreases in circularity (Figure 4D; average change of 0.5%; median, -5.8%). These data suggest that contractility is required specifically during compaction to establish niche architecture.

**Figure 4:**
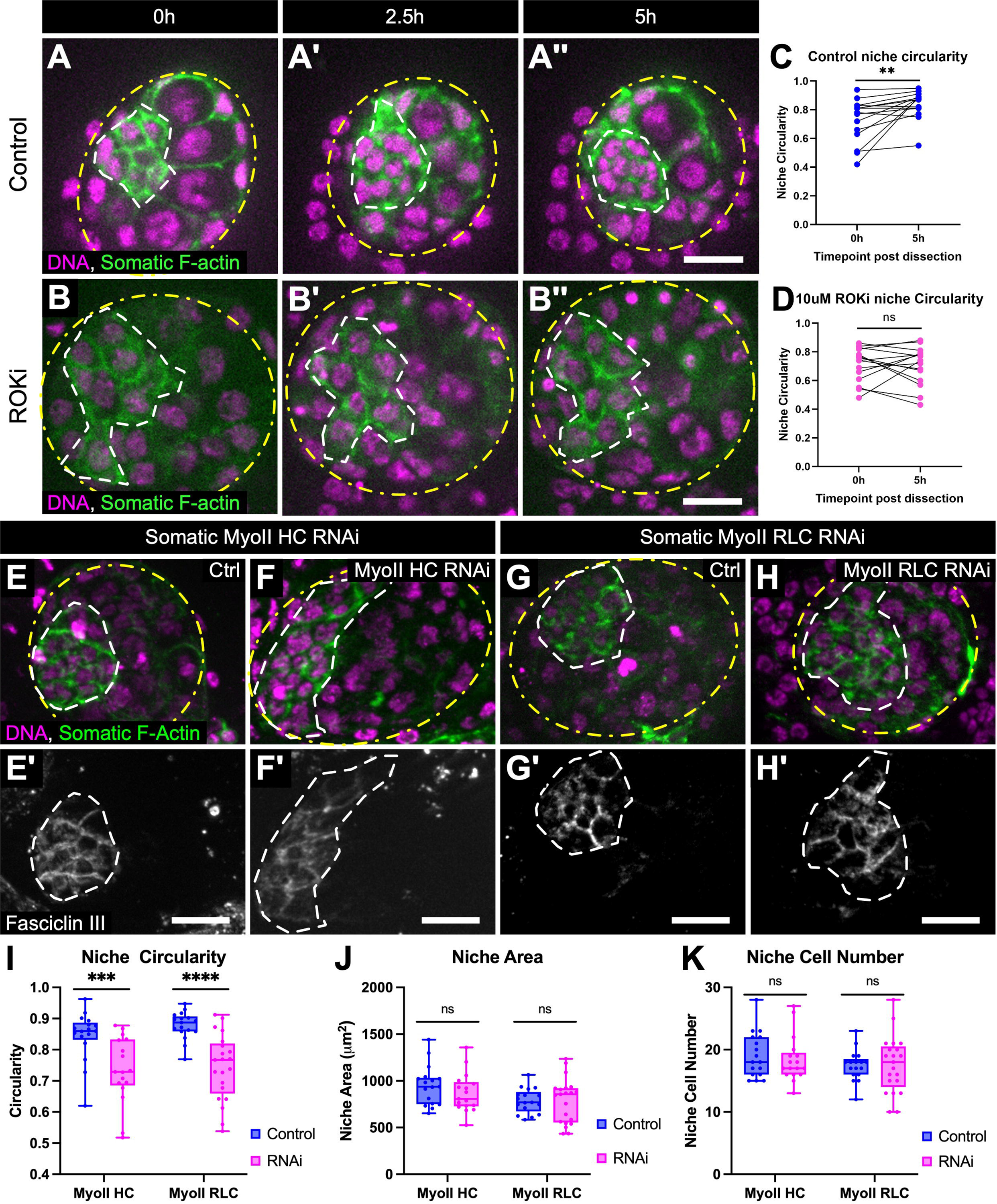
AMC is required for niche morphogenesis. A-B) 5-hour timelapse of control (A) or ROKi-treated (B) Six4-Moe::GFP (somatic F-actin, green); His2Av::mRFP1 (nuclei, Magenta). C-D) Niche circularity measurements at 0h and 5h of untreated (C, n=15 gonads; previously shown in ref.^12^) or ROKi-treated gonads (D, n=15 gonads). Lines pair the same niche at 0 and 5h (**p<0.01, Wilcoxon test). E-H) Six4-moe::GFP, His2Av::mRFP1 control (E, G) or Six4-Gal4 > MyoII RNAi gonads (F = HC RNAi; H = RLC RNAi) developed *in vivo.* E’-H’) FasIII. I) Niche circularity is decreased when either MyoII is depleted (***p<0.001, ****p<0.0001, Mann-Whitney; See Figure S1). J-K) Niche surface area (J) and niche cell number (K) show no difference between RNAi treatments and respective controls (Mann-Whitney). For graphs I-K, n=17 gonads for all conditions, except n= 21 for MyoII RLC RNAi.

As niche cells contribute to MyoII enrichment at niche-GSC interfaces (Figure 2E), we used Six4-Gal4 to challenge MyoII specifically in somatic gonadal cells. Somatic RNAi against MyoII HC successfully depleted MyoII at the niche-GSC interface (Figure S1), and led to severe compaction defects, as evidenced by a jagged niche-GSC boundary (Figure 4E, E’, F, F’). Niche circularity, but not niche area or cell number, significantly decreases compared to controls, and we observed similar defects using MyoII RLC RNAi (Figure 4G-K). These results show that somatic AMC is required for niche shape.

### Proper niche shape limits access to self-renewal signals

The reproducibility of niche morphology suggests that shape contributes to function. Of the niche-derived signals maintaining stem cells in adult testes, only Upd acts at this early stage. Upd expressed from niche cells activates the JAK-STAT pathway in cells nearest the niche, leading to accumulation of STAT, which is routinely used as a reporter for pathway activation^27,28^.

In both sibling controls and somatic MyoII HC RNAi gonads, germ cells adjacent to the niche exhibited STAT enrichment compared to germ cells further away (Figure 5A, A’, B, B’; quantified in C), demonstrating that improperly shaped niches can signal. However, compared with controls, more germ cells contact misshapen niches (Figure S2A), and most of these germ cells are activated for STAT (Figure 5D). In fact, we noticed a significant increase in all germ cells of the gonad (Figure S2B), likely due to a larger stem cell pool. As a critical role for the niche is establishing an appropriate number of stem cells, these results suggest that niche structure is essential for proper function.

**Figure 5:**
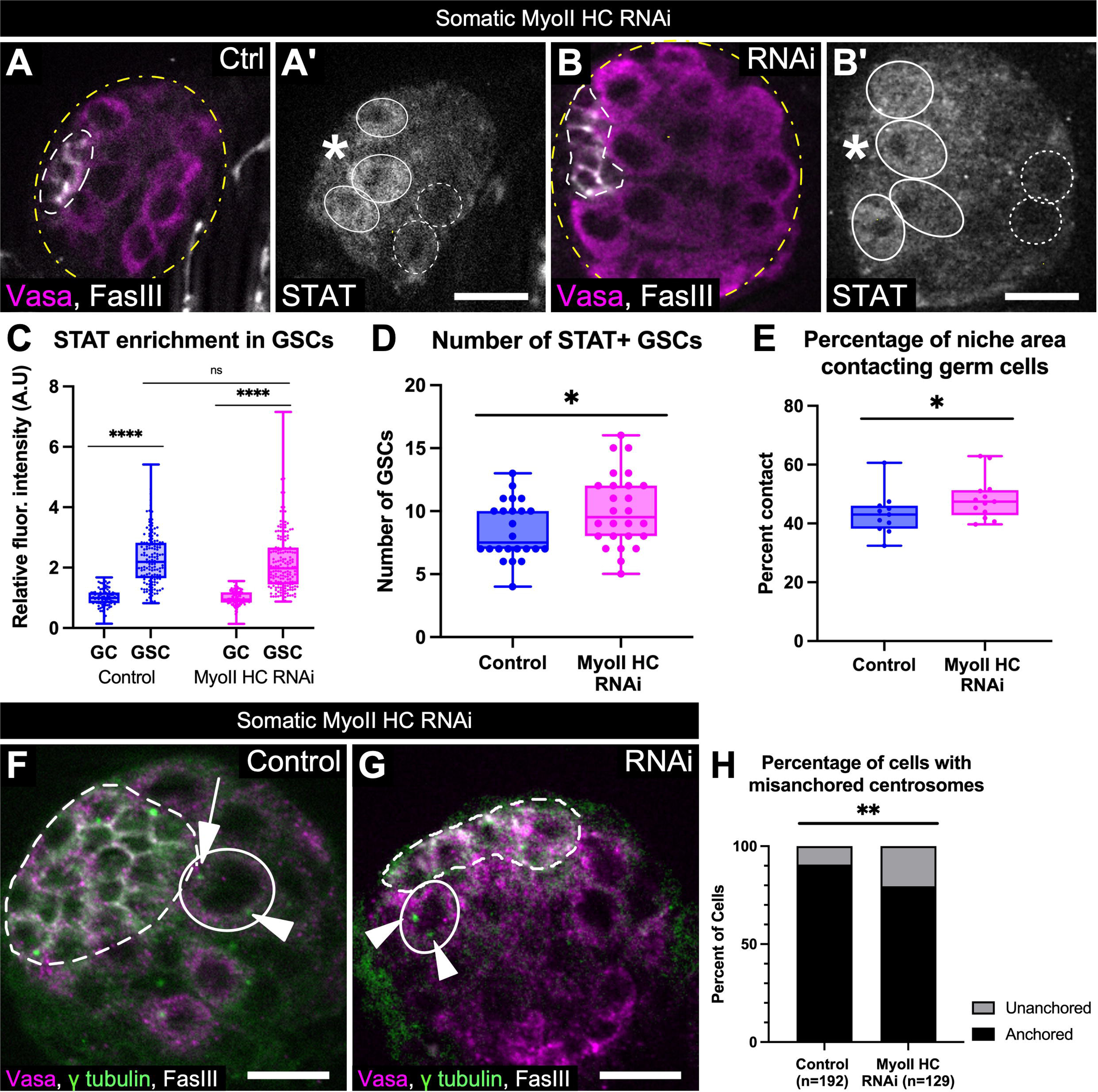
Proper niche shape is required to regulate GSC behavior. A-B) Control (A) or Six4-Gal4 > MyoII HC RNAi (B) stained gonads; Vasa (germline, magenta), FasIII (white). A’- B’) STAT antibody. Asterisk = niche. Solid outlines = niche-adjacent germ cells (‘GSC’) enriched for STAT. Dotted outlines = posterior germ cells (‘GC’) exhibiting lower STAT. C) STAT enrichment in GSCs under control (n= 87 GCs and 145 GSCs) and RNAi (n=108 GCs and 180 GSCs) conditions. D) Number of STAT+ germ cells contacting the niche is increased in MyoII HC RNAi gonads (n= 27 gonads) compared to controls (n=23 gonads). E) Percentage of niche area that contacts the germ line is increased in MyoII HC RNAi gonads. (*p<0.05, ***p<0.001, ****p<0.0001, Mann-Whitney; See Figure S2). F-G) Control (F) and MyoII HC RNAi (G) stained gonads; centrosomes (gamma tubulin, green), Vasa (germline, magenta), and FasIII (white). F) A GSC (solid outline) with one centrosome oriented at the niche (arrow), and one opposite (arrowhead). G) A GSC (solid outline) with two centrosomes (arrowheads) both oriented away from the niche. H) Percentage of cells with mispositioned centrosomes doubled from 9.4 to 20.5% comparing control and MyoII HC RNAi (**p<0.01, Chi-Square; See Figure S1 and S3).

Intriguingly, MyoII HC RNAi niches are not larger (Figure 4J), yet they support more stem cells. We hypothesized this might be explained by increased niche surface area accessible to germline. Typically, only a portion of the niche contacts the germline. We measured the percentage of niche surface physically contacting germ cells (Figure S2C, C’, D, D’), and indeed found an increase under MyoII HC RNAi conditions (Figure 5E). These data suggest that a compact niche is required to limit available surface area to restrict the number of GSCs.

### Proper niche shape is required for GSC centrosome orientation

Another critical niche function is to orient GSC divisions to ensure one daughter will remain a GSC near the niche, and the other will be moved away and differentiate^21–23,25,26^. To assess the role of niche shape on division orientation, we live-imaged control and ROKi-treated gonads, since this treatment affects niche shape (Figure 4A-D). ROKi will also arrest cytokinesis^37^, and indeed we captured completed GSC cytokinesis in one third of the control gonads (11/32), but never in ROKi-treated gonads (0/26). Whether cytokinesis completed or not, we could map the orientation of nuclear divisions using His2Av::mRFP1 to calculate chromatin trajectory relative to the niche-GSC interface (Figure S3A). In these analyses, higher angles represent divisions oriented away from the niche^38^. Control nuclei divided predominantly from 30-80 degrees (Figure S3B, B’, D, E), indicating that GSC division orientation is regulated and biased so daughter cells are born further from the niche. In contrast, division orientation of ROKi-treated cells significantly differed from controls, with GSCs primarily dividing between 0-45 degrees relative to the niche (Figure S3C, C’, D; p<0.001, KS test; Figure S3F), which is consistent with a randomized distribution with 3-dimensional constraints taken into consideration^38^.

A limitation is that pharmacological treatment affects all cells. To determine whether changes in niche shape affect GSC behavior non-autonomously, we manipulated niche shape in a lineage-specific manner. Division orientation is controlled via anchoring of one GSC centrosome at the niche-GSC interface^21–23,25,26^. We therefore analyzed centrosome positioning in fixed tissue after somatic MyoII knockdown. In controls, as expected, about 90% of GSCs had proper centrosome positioning (Figure 5F, H). In contrast, the frequency of mispositioned centrosomes doubled in GSCs from gonads exhibiting somatic MyoII knockdown (Figure 5G-H). By chance movement, unanchored centrosomes abut the niche ∼50% of the time^23^; thus, change of this magnitude is significant.

Taken together, niche shape is crucial in regulating GSC centrosome orientation and in limiting self-renewal signals to a subset of cells.

### GSCs are required to shape their niche

We sought to uncover mechanisms contributing to actomyosin polarity within the niche required for proper form and function. Importantly, GSCs are recruited to the niche during morphogenesis^27^, and this coincides with GSC division onset^12^. Since mechanical forces can polarize MyoII^39,40^, a potential force on niche cells could be the packing or cell divisions that occur as GSCs are recruited to the niche.

Evidence that forces emanate from GSCs first came from laser-severing experiments carried out during early compaction, prior to MyoII polarization (Figure 6). When tension along the niche-GSC interface was severed, the germ cell protruded into the niche (Figure 6A, A’, B), suggesting exertion of positive pressure on the niche. Interestingly, this force appears exclusive to early compaction, as no protrusion was observed when cuts were performed nearing the end of compaction (Figure 6C, C’, D). The timing of the force exerted by germ cells on the niche, and the eventual polarization of MyoII along the niche-GSC interface, led us to hypothesize that germ cell force contributes to polarization required for niche shape.

**Figure 6:**
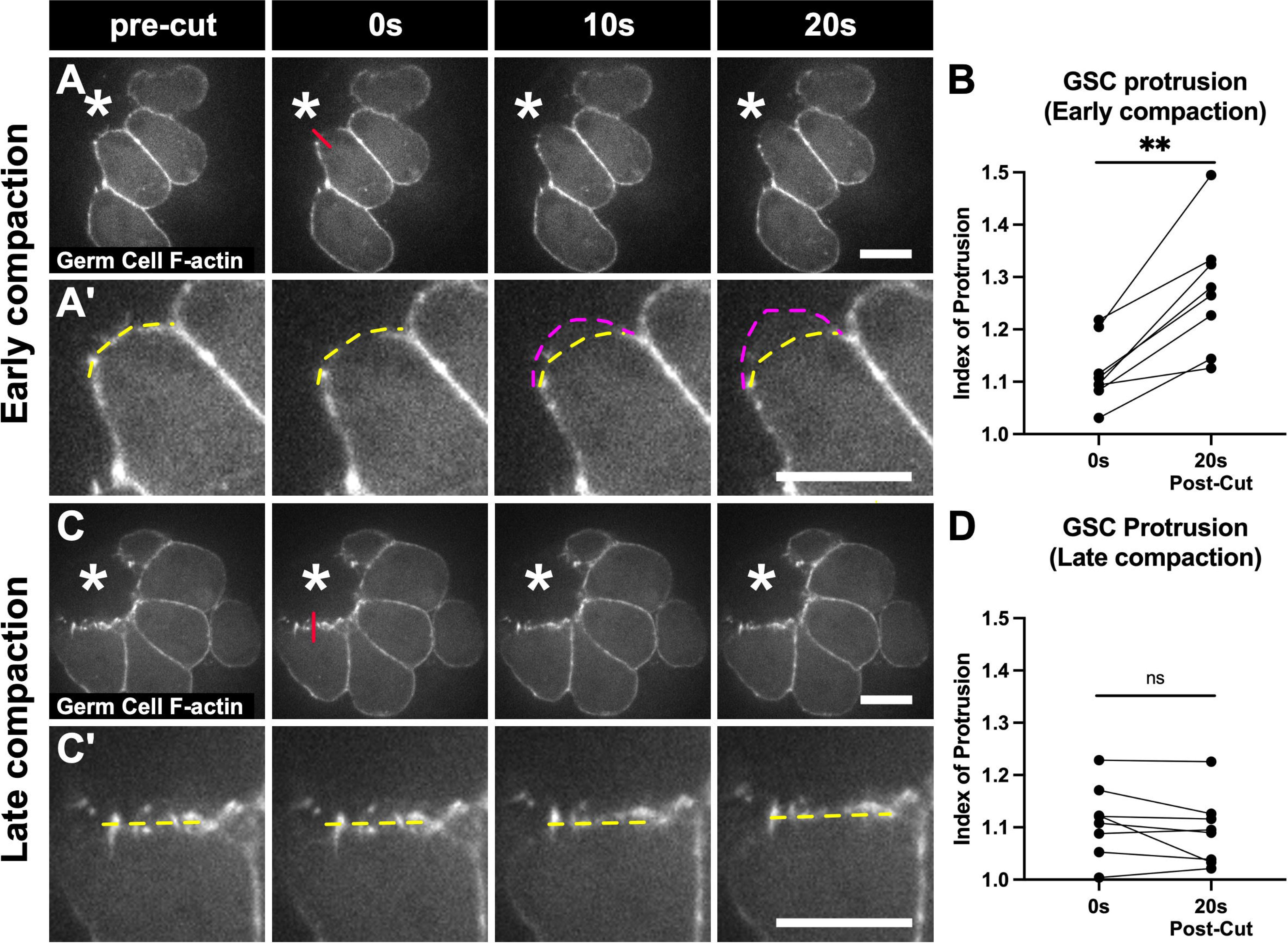
Germ cells protrude into the niche during early compaction. A, C) Gonads *ex vivo* expressing Nanos-Moe::GFP (germline F-actin); early (A) or late compaction (C). Asterisk = niche. Niche-GSC interface contour was monitored before laser cut (pre-cut), during cut (red line, 0s), and 10 and 20s post-cut. A’, C’) Inset of analyzed cell. Yellow dashed line = contour pre-cut; magenta = extent of protrusion post-cut, if any. A, A’) Severing the niche-GSC interface during early compaction led to germ cell protrusion into the niche (10s and 20s). C, C’) Severing the niche-GSC interface during late compaction revealed little to no protrusion. B, D) Quantifications of protrusion into the niche, where each line represents the same interface 0s and 20s after severing. Germ cells significantly protrude into the niche during early (B) but not late compaction (D; n=8 cells each; **p<0.01, Wilcoxon Test).

We therefore selectively ablated germ cells by expressing the pro-apoptosis gene, *head involution defective* (*Hid*), using Nanos-Gal4::VP16 (Figure 7A-B). We found that upon Hid expression, gonads contained fewer germ cells, confirming ablation (Figure 7E), and had misshapen niches (Figure 7B, F).

**Figure 7:**
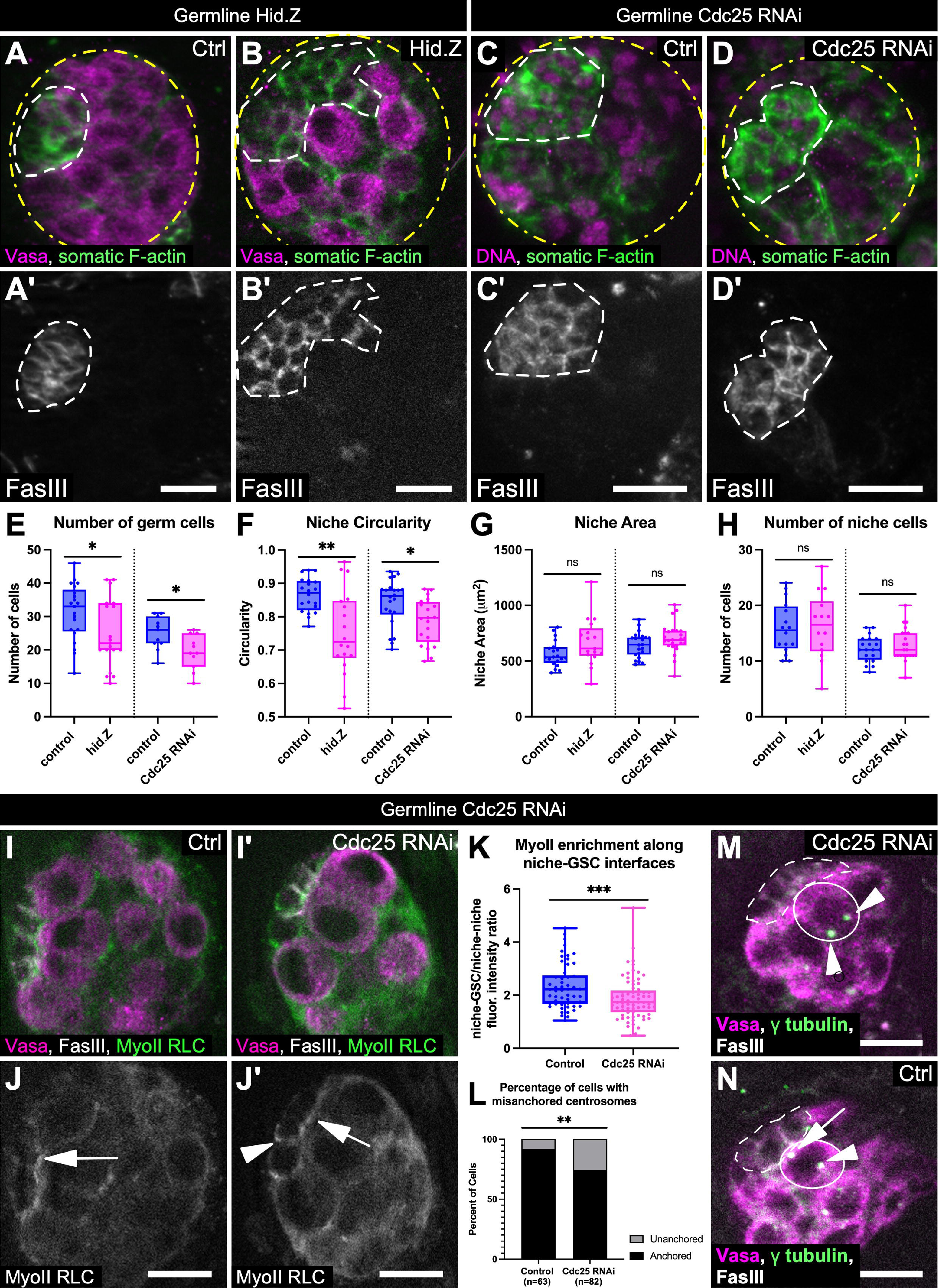
GSC divisions are required to shape their niche. A-B) Six4-Moe::GFP control (A) or Nanos-Gal4::VP16 > hid.Z gonad (B) developed *in vivo*; F-actin (green), Vasa (magenta), and FasIII (A’-B’). C-D) Six4-Moe::GFP; His2AV::mRFP1 control gonad (C) or Nanos-Gal4::VP16 > Cdc25 RNAi gonad (D) developed *in vivo*; F-actin (green), DNA (magenta) and FasIII (C’-D’). A, C, and D are maximum projections of ∼10 slices (0.5-micron intervals). E) Total germ cell number is reduced by expressing hid.Z (n=19 gonads) or Cdc25 RNAi (n= 11 gonads) in the germline compared to respective controls (n=21 and 11, respectively). F) Niche circularity is decreased upon germline expression of hid.Z or Cdc25 RNAi (*p<0.05, **p<0.01, Mann-Whitney). G-H) No difference in niche surface area (G) nor niche cell number (H) between conditions (n=18 hid.Z and 21 Cdc25RNAi gonads) and respective controls (n= 20 and 22, respectively). I-J) Control (I) and Cdc25 RNAi (J) gonads expressing MyoII RLC::GFP; Vasa (magenta), FasIII (white). I’-J’) MyoII RLC::GFP shows enrichment in controls towards niche-GSC interfaces (I’, arrow), and decreased but residual polarity towards the niche-GSC interface in Cdc25 RNAi gonads (J’, arrow). MyoII is sometimes mis-polarized away from the niche-GSC interface (J’, arrowhead). K) Quantifications show germline Cdc25 RNAi leads to decreased MyoII enrichment at the niche-GSC interface (n=54 control = interfaces, n=72 Cdc25 RNAi interfaces, ***p<0.001, Mann-Whitney; see Figure S4). L) Percentage of cells with mispositioned centrosomes tripled from 8 to 26% comparing control and Cdc25 RNAi gonads (**p<0.01, Chi-Square). M-N) Control (M) and Cdc25 RNAi (N) gonads; gamma tubulin (green), Vasa (magenta), FasIII (white). M) A GSC (solid outline) with one centrosome anchored at the niche (arrow), and one located opposite (arrowhead). N) A GSC (solid outline) with two centrosomes (arrowheads) oriented away from the niche.

We suspect that the latent protrusive force exerted by germ cells (Figure 6A, B) has various origins. One contribution might be from mitotic divisions^41^. We compromised germ cell divisions by lineage-specific knock down of Cdc25^42^ via RNAi. Upon Cdc25 knockdown, germ cell number decreased (Figure 7E), confirming divisions were inhibited, and observed defects in niche circularity (Figure 7C, D, F). Neither niche area nor niche cell number changed upon GSC division inhibition or germ cell ablation (Figure 7G, H), which parallels our findings from MyoII knockdown in the niche (Figure 4J, K). These data confirm that GSCs act to shape their niche and suggest that GSC divisions at least partially contribute to the mechanical force required to establish appropriate niche morphology.

### GSC divisions aid in polarizing MyoII within the niche

We have established that MyoII polarity is essential for niche architecture. To assess whether germline divisions regulate this polarity, we visualized MyoII RLC::GFP in Cdc25 RNAi and sibling controls. In controls, MyoII was enriched along the niche-GSC interfaces (Figure 7 I, I’, K), but this was significantly decreased in Cdc25 RNAi gonads (Figure 7J, J’, K). Notably, some niche cells polarized MyoII away from the niche-GSC boundary (Figure 7J’, arrowhead). Further, Cdc25 RNAi gonads exhibited an increase in centrosome mispositioning, suggesting that these niches have functional defects (Figure 7L-N). Thus, GSC divisions are required to shape their niche by polarizing MyoII to the stem cell-niche interface, which is critical for niche function.

Adherens junctions (AJs) are a main source of niche-GSC adhesion^26,45^ and can control mechanosensation in other contexts^43,44^. To test whether cadherins could be the sensor for germline forces on niche cells, we knocked down E-cadherin, but observed no defect in niche shape (Figure S4). Thus, factors underlying mechanotransduction remain to be discovered in this system.

Collectively, however, the effects we observed on niche shape and function by depleting germ cells or inhibiting their division strongly suggest that GSCs feedback to shape the niche that guides their behavior.

## Discussion

Our work captures the dynamic formation of a functional niche. Prior imaging revealed how pro-niche cells migrate to their proper position^12,30^. Here, we show that its continued morphogenesis is dependent on polarization of a force-producing actomyosin network. If contractility is compromised, the niche adopts an irregular contour and exhibits defects in function. Additionally, recruited stem cells aid in polarizing MyoII along the forming niche-stem cell boundary. Without this feedback, niche shape and function are aberrant.

### Polarized contractility is required for niche morphogenesis

Cytoskeletal organization is crucial for compacting niche cells into their functional arrangement. Specifically, F-actin and MyoII induce tension along the interface between niche cells and newly recruited GSCs during compaction. Compromising contractility yielded misshapen niches, reminiscent of the involvement of MyoII for mammalian intestinal niche curvature^7^, intestinal stem cell shape^8^, and neural stem cell rosette morphogenesis^46^. Other studies benefitted from whole-tissue knockouts, pharmacological manipulations, and/or *in vitro* culturing, but were limited in cell-type specificity. Our study combines *ex vivo* live-imaging^29^ with *in vivo* cell-type specificity. While pharmacological manipulation affects all cells, the asymmetry in MyoII contribution suggests that it is the contractile activity in the niche cell cortex that is a main driver in niche morphogenesis. This inference was strongly supported by the misshaping observed when MyoII was depleted only in somatic cells.

It would be interesting to investigate whether AMC in the GSC cortex also shapes the niche, but this is not currently testable^47^. Indeed, our ROKi experiments support such a conclusion. Tissue-level inhibition of AMC induced more severe phenotypes than somatic MyoII inhibition (Figure 4C, F, H). Of note, ROKi treatment *both* altered MyoII function in the niche *and* disrupted GSC cytokinesis. We suspect the severe niche phenotype of ROKi treated gonads is caused by a combinatorial effect of GSC and niche cell involvement in niche morphogenesis.

### Niche morphology regulates niche-stem cell signaling

Our data suggest that the contour of the niche is important for modulating signaling. We previously reported that blocking niche assembly led to niche cells that signaled poorly, if at all^30^. By affecting compaction, we find significant STAT activation in nearby germ cells, but an increase in the number of those cells contacting the niche. It is possible that MyoII inhibition increases Upd secretion from niche cells. However, because the pathway receptor polarizes along the niche-GSC interface within GSCs^48^, we believe the increase of niche surface available to the germline likely accounts for the increase in niche-adjacent STAT-positive cells. Work in intestinal organoids has also shown that niche curvature helps optimize signaling^8^, highlighting that concepts uncovered in *Drosophila* are applicable to mammals.

### Centrosome orientation is dependent on a precise niche shape

Gonads with aberrant niche morphology exhibit defects in centrosome positioning. Normally, the mother centrosome anchors to the GSC cortex at the interface with the niche via influence of proteins polarized to this interface^21,25,26,48^. If centrosome anchoring fails, a “centrosome orientation checkpoint” (COC) is engaged, preventing spindle assembly to avoid symmetric divisions^22,23^.

Prior centrosome orientation research has exclusively manipulated factors acting in the germline cortex^21,25,26,48^. Our work is the first to reveal a requirement on the niche side of this interface impacting centrosome positioning. An intriguing model is that polarized MyoII-dependent contractility in the niche is required to polarize yet-to-be-identified, novel COC players to the niche cortex.

### GSCs feedback to shape their own niche by polarizing MyoII

Importantly, germ cells are required for proper niche shape and function. Specifically, GSC divisions are required for niche MyoII polarization, and consequently proper niche shape and function. We speculate there is a mechanosensitive pathway between dividing GSCs and the niche, perhaps activated by force-producing spindle microtubules on the GSC cortex^41^. It is also possible that a cell cycle-dependent change to GSC polarization causes the effect on niche cell polarity.

Future work must discern how the niche senses force to elicit a response in cytoskeletal polarity. While E-Cadherin is not essential (Figure S4), perhaps a combination of AJ components^49^ must be inhibited to affect niche morphogenesis. Alternatively, other mechanisms, such as mechanosensory ion channels,^50^ may be involved.

Additionally, division forces are likely not the only source of force from the germline. Protrusion into the niche upon cortical laser severing suggests there is latent germline pressure toward the niche. Although we do not know the mechanistic basis for that pressure, the residual MyoII polarization when germline divisions are compromised supports the idea that additional forces generated by GSCs contribute to niche morphogenesis. Indeed, removing many of the germline cells by inducing cell death led to more severe niche phenotypes than inhibiting divisions (Figure 7F, compare circularity changes).

Feedback between stem cells and their niche has been seen in other systems, such as the *C. elegans* hermaphroditic gonadal niche^51^, the murine hair follicle stem cell niche^52,53^, and the *Drosophila* glial niche^54^. Therefore, our work strengthens the paradigm that niche-stem cell relationships are bidirectional and identifies mechanisms of stem cell force-generation to regulate niche form and function.

## Limitations

While our lineage-specific manipulations show somatic requirement for contractility in niche compaction. The only niche driver available for these stages, Six4-Gal4, is also expressed in cyst cells. Thus, determining whether cyst cells also impact niche shape in a MyoII-dependent manner is not yet possible. Cyst cells make much less extensive connections with the niche as do germ cells (c.f. Figure 2A’; the relative thinness of somatic-to-niche contacts is apparent) ^17,33^. We therefore believe it unlikely that cyst cells directly affect niche morphogenesis. Nevertheless, cyst cells could signal or provide force to the GSCs as they shape the niche.

Additionally, Hedgehog and Bone Morphogenetic Protein (BMP) are produced by the adult niche, but a role for neither has been described during niche formation. Hedgehog acts exclusively on CySCs, which are recruited later in larval stages^28,55^. In the adult, BMP signaling requires intimate contact between GSCs and niche cells^45,56,57^. While it would be interesting to determine how niche shape impacts this pathway, activation is only robust enough in later larval GSCs for analysis^58^.

Lastly, we are curious as to whether mechanisms revealed in building this niche are conserved at steady state. Preliminary work on the adult testis suggests enrichment for contractility regulators in the niche. Therefore, approaches available in the adult testis might provide answers to the above questions.

## Supporting information

Reagents Table

## Acknowledgements

We thank BDSC (NIH P40OD018537), the Lehmann, Bach, Martin, Bellaïche, and Kiehart labs and DSHB for reagents. We thank K. Ong and A. Stout for advice on laser cutting. Thanks to G. Vida and the Lenhart lab for input, K. Lenhart for revision assistance, and R. Mudgett-McGeoch for coding assistance. This work was supported by NIH: F31HD105342, T32GM007229 to BNW; T32HD083185, F31HD111208 to KAN; GM125123 to LA; R35GM136270 to SD.

## Author Contributions

BNW: Methodology; Validation; Formal Analysis; Investigation; Data Curation; Writing-Original Draft; Writing-Review-Editing; Visualization; Funding Acquisition. KAN: Methodology; Software; Validation; Formal Analysis; Investigation; Data Curation; Writing-Review-Editing; Visualization. JS: Methodology; Formal Analysis; Investigation; Data Curation; Visualization. LA: Methodology; Formal Analysis; Writing- Review & Editing; Supervision. SD: Conceptualization; Methodology; Validation; Investigation; Resources; Writing-Review-Editing; Visualization; Supervision; Project Administration; Funding Acquisition.

## STAR Methods

### RESOURCE AVAILABILITY

#### Lead Contact

Further information and requests for resources and reagents should be directed to and will be fulfilled by the lead contact, Stephen DiNardo.

#### Materials Availability

Six4-Gal4::VP16 *Drosophila* stocks will be provided upon request to the lead contact.

#### Data and Code Availability

All data reported in this paper will be shared by the lead contact upon request. This paper does not report original code. Any additional information required to reanalyze the data reported in this paper is available from the lead contact upon request.

### EXPERIMENTAL MODEL AND SUBJECT DETAILS

#### Transgenic Fly stocks

##### Gal4 Stocks

Nanos-Gal4::VP16 was used to restrict expression of UAS-responder transgenes to germline cells^59^. To restrict expression to somatic gonadal cells, we utilized Six4-Gal4 and Six4-Gal4::VP16 transgenic lines. For knockdown experiments, we utilized the previously described Six4-Gal4^12^, aging embryos for 15-17 hours at 29 degrees C to increase knockdown efficiency, capitalizing on the inherent temperature sensitivity of full length Gal4. To label cytoskeletal proteins with cell-type specificity, we generated a stronger line where the Gal4 DNA binding domain was fused to a VP16 transactivation domain (Six4-Gal4::VP16).

To construct the Six4-Gal4::VP16 line, a previously identified enhancer from the third intron of Six4^60^ was amplified from genomic DNA with primers listed in the Reagent Table, and cloned into the pENTR/D-Topo entry vector (Invitrogen; K240020). The enhancer was transferred into the destination vector pBPGal4.2::VP16Uw (ref.^61^; Addgene, # 26228 ; RRID:Addgene_26228) via an LR clonase reaction. Transgenic flies were generated by The Best Gene, using PhiC-31 integration into an attP host (RRID:BDSC_8622).

##### Fluorescent labeling

For cytoskeletal or nuclear labeling, we utilized multiple fluorescent transgenic constructs. Nuclear chromatin was labeled using a His2Av::mRFP1 transgene (FBtp0056035). Somatic F-actin was labeled by P{D-six4- egfp:moesin (here called Six4-Moe::GFP; ref.^62^) or Six4-Gal4 > UASp-F-Tractin::TdTomato (a recombinant of Six4-Gal4 and RRID:BDSC_58989), whereas Germ cell F-actin was labeled by Nanos-Moesin::GFP^63^. To visualize MyoII with lineage specificity, we utilized a UAS-GFP::Zipper construct^34^ driven by either a somatic or germline Gal4. Ubiquitous MyoII was visualized using either sqh-Sqh::mCherry (ref.^31^; RRID:BDSC_99923; RRID:BDSC_59024) or sqh-Sqh::GFP3x^64^.

##### Transgenic manipulation

The following transgenes expressing shRNAs were used in knock down experiments: MyoII HC (RRID:BDSC_65947), MyoII RLC (RRID:BDSC_33892), Cdc25 (RRID:BDSC_34831), and E-Cadherin (RRID:BDSC_32904). P(UAS-hid.Z)2 (RRID:BDSC_65403) was used to ablate germline cells. Knockdown of the MyoII heavy chain was confirmed by measuring fluorescence intensity of Sqh-Sqh::GFP3x along the niche-GSC interface normalized to unmanipulated GSC interfaces (see cytoskeletal polarity analysis section for more details). Germ cell ablation or division inhibition were confirmed by quantifying the number of germ cells in manipulated gonads compared to sibling controls and analyzed via a Mann-Whitney test.

In MyoII HC RNAi experiments, some gonads were defective in niche assembly, as evidenced by multiple niche aggregates. This is likely because timing of the somatic Gal4 driver cannot be controlled as accurately as, for example, our pharmacological ROKi treatment. Gonads exhibiting niche assembly defects were excluded from analysis.

##### Fluorescent balancers

For all transgenic manipulation experiments, utilization of marked balancers allowed unambiguous identification of embryonic gonads that expressed both the Gal4 and UAS construct, as well as sibling controls that lacked either the Gal4 or the UAS construct, as identified by the presence of the appropriate fluorescent balancer (TM3, P{w[+mC]=Gal4-twi.G}2.3, P{UAS-2xEGFP} AH2.3, Sb[1], Ser[1], FBst0006663; TM6B, P{Dfd- GMR-nvYFP}4, Sb[1] Tb[1] ca[1], RRID:BDSC_23232; or CyO, P{w[BmC]=Dfd-EYFP.w[BmC]}2, RRID:BDSC_8623).

For simplicity, we refer to the latter two balancers as DfdGFP.

#### Sex identification

Male embryos, as well as gonads dissected from male embryos, were identified by the presence of male-specific, msSGPs, visualized by high-level expression of the Six4-Moe::GFP transgene, or by the accumulation of Vasa protein as previously described^12^.

### METHOD DETAILS

#### Fixed gonad preparation and staining

##### Embryonic gonad dissections and Fixation

Unless otherwise noted, embryos were collected and aged in a humid 25 degree C chamber. Embryos were either aged 15-17 hours after egg lay (ES16) for pre-compaction niche analysis or 22-24 hours (late ES17) for post-compaction niche analysis. Embryos were dechorionated, staged based on gut morphology, hand- devitellinized and dissected in ∼500uL of Ringers solution (5mM HEPES, pH 7.3; 130mM NaCl; 5mM KCl; 2mM MgCl_2_; 2mM CaCl) as previously described ^12^. Tissue was fixed in 4% formaldehyde, Buffer B (16.7mM KPO, pH 6.8; 75mM KCl; 25mM NaCl; 3.3mM MgCl_2_), and 0.1% Triton-X-100 for 15 minutes, then washed in PBS (10mM Na_2_ HPO_4_; 1.8mM KH_2_PO_4_; 2.7mM KCl; 137mM NaCl; pH 7.4) plus 0.1% Triton-X-100 (PBS-Tx). Tissue was then blocked for 1 hour at room temperature in 4% normal donkey serum in PBSTx.

##### Immunostaining

Primary antibodies were used overnight at 4C. We used Goat antibody against Vasa 1:400 (Santa Cruz, dC-13, now discontinued); Rabbit antibody against STAT92E 1:350 (gift from E. Bach, NYU); and RFP 1:1000 (Abcam, ab62341); mouse antibody against Fasciclin III 1:50 (DSHB, 7G10); and Gamma Tubulin 1:200 (Sigma, GTU-88); Rat antibody against E-Cadherin (DSHB, DCAD2); and chick antibody against GFP 1:1000 (Aves Labs, GFP-1020). Secondary antibodies were used at 3-4ug/ml (Alexa488, Cy3, orAlexa647; Molecular Probes and Jackson ImmunoResearch) for 1 hour at room temperature. DNA was stained with Hoechst 33342 (Sigma) at 0.2ug/ml for 5 minutes. Tissue was equilibrated overnight in 50% glycerol and 50% Ringers, then mounted with 2% propyl-gallate in 80% glycerol. Images of fixed gonads were acquired on a Zeiss Imager with Apotome using a 40x, 1.2 N.A. lens.

#### Ex-vivo experiments

##### Live-imaging

Dissection and live-imaging was performed as previously described^29^. Embryos were aged in a humid container in a 25-degree incubator for 14-17 hours after egg lay, and staged based on gut morphology to select stage 16 embryos. For analysis of gonads before and after compaction, we imaged for 5 hours. We only imaged gonads with niches that have been assembled at the gonad anterior but the contours of which have not yet become rounded. For some experiments, we selected gonads based on their stage within compaction (early, mid, or late) depending on the relative smoothness of the niche boundary. Imaging was carried out on gonads carrying either a somatic (Six4-Moe::GFP) or germline (Nanos-Moe::GFP) F-actin marker, and in some cases a MyoII marker (sqh-Sqh::mCherry). Gonads were imaged using an IX7 Olympus spinning disk confocal, using a 63x, NA 1.2 water immersion, or a 100x, NA 1.4, oil immersion lens, and captured with an EMCCD camera (Hamamatsu photonics, model C9100-13) controlled by MetaMorph software.

##### Laser ablation

Laser ablation experiments were carried out on dissected gonads carrying either a somatic (Six4-Moe::GFP) or germline (Nanos-Moe::GFP) marker. We prepared gonads for live-imaging as described above. However, we imaged gonads at two different stages of compaction: early or late. Identification of niches undergoing early compaction is described above. To analyze late-compaction stages, such as in Figures 3 and 6, we again selected stage 16 embryos, but only those whose niches were both assembled and rounded.

To identify prospective interfaces to be targeted for ablation, a single time point z-stack was acquired for each gonad. Usually only one or two interfaces would be selected for treatment per gonad to limit any potential effects of global relaxation. The ablating beam generated by a MicroPoint laser emanating from a 405 nm dye cell was focused to the interface through a 100x, 1.4NA lens, using Andor IQ3.2 software. The micropoint laser settings were optimized at each session, selecting the minimum power required for junction severing.

Simultaneously, Metamorph software was set to stream single color (488 excitation) images from the plane of the interface at 250 milliseconds intervals. Streaming acquisition was begun, and then the laser fired to treat that interface selectively. Post-treatment acquisition would continue for 1–3 minutes.

##### Rho Kinase Inhibitor treatment

For inhibitor treatments, a single timepoint z-stack was obtained for each gonad to establish a pretreatment standard. Then, the potent and selective Rho Kinase inhibitor, H-1152, was added to 10 μM final concentration with thorough but gentle pipetting (Santa Cruz, sc-203592; Ki = 1.6 nM for Rho Kinase compared with Ki = 140 nM for the inhibitor Y-27632). Time-lapse imaging was carried out for 5 hours, with inhibitor replenished every 90 minutes^65^. In experiments where laser ablation of the cortical cytoskeleton was carried out, the ablations were begun within 15-30 minutes after inhibitor addition.

### QUANTIFICATION AND STATISTICAL ANALYSIS

Fixed Images were analyzed via the Image J Blind Analysis Tools plugin.

#### Cytoskeletal polarity analysis

Fluorescent gonads were dissected and immunostained either with an antibody against GFP or RFP, depending on the fluorescent transgene present. Niche interfaces were visualized by anti FasIII. To extract the fluorescence intensity of F-actin or MyoII, a 3-pixel wide line was drawn over the target niche-GSC interface and its mean grey value returned in Image J. In most cases, the control interfaces were, respectively, either niche-niche or GSC-GSC interfaces depending on whether niche cells or germ cells were being analyzed. While confirming MyoII knockdown, however, GSC-GSC interfaces were utilized for normalization, since these interfaces should not have been affected by MyoII knockdown. Mean gray values were background subtracted by drawing a line where no tissue was present, and niche-GSC interface values were normalized to an average of the respective control interfaces taken for that gonad. Normalized values were plotted, and niche-GSC intensity values were either compared between genotypes, or compared to control interfaces of the same genotype, using a Mann-Whitney test.

#### Niche phenotypic analysis

For analysis of niches from *ex vivo* cultured gonads, niche circularity was measured as described previously ^12^. For fixed samples, three dimensional images were displayed using Imaris software. Niche area was measured by generating two surfaces in Imaris. The first surface was manually created by using FasIII and Six4-Moe::GFP as a guide to outline the niche on multiple z-planes. Creating this first surface is essential to isolate the niche from surrounding tissue. However, the manually drawn surface is a rough outline of the niche, and therefore needed to be refined to accurately recapitulate the curvature of the niche. To make a more refined surface, we made a mask of Six4-moe::GFP fluorescence from the first, rough surface. Using the masked Six4-moe::GFP fluorescence, we generated the second surface via Imaris’s automatic surface generation protocol. We smoothed the surface with a 0.5-micron surface detail, and pixel-value thresholds were determined manually to ensure that the entire niche boundary was included in the surface. Niche surface area measurements were extracted from the second, refined surface for all samples by using the Imaris vantage function.

To measure the percentage of niche area that contacted germ cells, we additionally created a surface of all germ cells in the same way we made our niche surface (see above), but instead used Vasa as our marker to generate the surface. We used a surface-surface contact Xtension to generate the percentage of the niche surface that was contacting the germline surface.

Niche cell counts were extracted in Imaris using either Hoechst or His2Av::mRFP1 to visualize individual cell nuclei and FasIII to visualize cell outlines. Nuclei were also marked to extract internuclear distance between each cell, and generate the average nuclear distance between a niche cell and its 3 nearest neighbors. Nuclear distance between the 3 nearest neighbors were extracted from all samples using the Imaris vantage function.

Finally, the 3-dimensional image was rotated to orient the niche head-on, and a screenshot was captured to measure circularity using Image J. To measure circularity, an ROI was drawn using the freehand selection tool to trace the niche boundary labeled by Six4-moe::GFP, and circularity was extracted using the ‘shape descriptors’ tool.

Mann-Whitney tests were used to compare niche parameters between different genotypes. A Wilcoxon-test was used for paired analysis when measuring niche circularity of gonads live-imaged *ex vivo* at two separate timepoints.

#### Quantification of retraction velocity

After live-acquisition, images were imported into Image J, with the treated interface identified, oriented vertically, and the time stack cropped. A montage was then created that consisted of one pre-treatment frame, followed by 60 seconds worth of post-ablation frames, with each frame in the montage at 5 second intervals; analyses carried out at 1 second intervals generated the same results. Using the segmented line tool, the junction above and below the cut interface of the niche cell and germline cell was marked throughout the montage. The X-Y coordinates for the mapped vertices were exported to a spreadsheet. Peak retraction velocities were determined from vertical displacement over time, as presented in the respective scatter plots. Peak retraction velocities between treatment conditions or interfaces were compared via a Mann-Whitney test.

#### GSC protrusion quantification

In those cases where severing the cortical cytoskeleton led to pronounced invasion of niche cell territory by the germ cell, the extent of invasion was quantified in the Nanos-Moe::GFP background by measuring the length of the protrusion from its base by tracing the arcing germ cell outline along the protruded part and comparing that to the vertical distance at the base of that protrusion. In this manner, no or little protrusion would yield a value close to one, while protrusions would yield values greater than one. Level of protrusion was compared between GSCs at early and late compaction timepoints with a Wilcoxon test.

#### Quantification of STAT accumulation

STAT intensity was extracted in Image J from anti STAT-stained gonads using regions of interest (ROI) drawn to include the Vasa signal of germline cells. In Figure 5C, for each gonad, we sampled 5 GSCs defined as those germ cells in contact with the niche, and 3 posterior germ cells located more than one cell diameter from the niche. We background-subtracted the average fluorescence intensity from each GSC and GC by drawing an ROI directly outside of the gonad. Relative STAT enrichment values were obtained by dividing the background-subtracted value of each GSC or GC by an average of the 3 background-subtracted GC values from that particular gonad. For Figure 5D, STAT positive cells were quantified by measuring the normalized fluorescence intensity of each germ cell directly contacting the niche as described. To establish a threshold over which a cell would be scored as STAT-positive, we took the average STAT enrichment (2.29 fold) and standard deviation (0.85) of all control cases. We then counted a cell touching the niche as positive if STAT levels were greater than 1.44-fold enriched (average enrichment minus 1 standard deviation). Data was obtained from sibling control and MyoII HC RNAi conditions and analyzed with a Mann-Whitney test.

#### Centrosome orientation analysis

Centrosomes were visualized with immunofluorescence against Gamma tubulin. GSCs contacting the niche were only scored for centrosome position if they had 2 clearly discernible centrosomes, and therefore had undergone centrosome duplication. For each such GSC, we scored whether one of the two centrosomes was located at the niche, as evidenced by cortical localization (visualized with Vasa) and proximity to the niche (visualized with FasIII). GSCs with one centrosome located along the niche-GSC interface were scored as “anchored,” whereas GSCs with neither centrosome located at the niche-GSC interface were scored as “unanchored.” Data was analyzed via Chi-squared analysis.

#### GSC division angle analysis

We surmised that the spatial constraints imposed by the spheroidal gonad might influence the possible division angles compared to that reported for the adult testis. That caution appeared justified, as using the rigorous 3-dimensional analysis described next, Figure 5K indeed showed that the distribution of angles was slightly broader than expected from previously published 2-dimensional centrosome and spindle analyses^21,25,66^. To preserve the 3-dimensional nature of the division coordinates, we used the model from ref.^38^. Therefore, we calculated division angles from live-imaged Six4-Moe::GFP;His2AV::mRFP1 gonads, based on two parameters as described by ref^38^: 1) a plane of best-fit to represent the niche interface, and 2) the division trajectory. First, a plane representing the niche-GSC interface was approximated as follows. The plane was highlighted by Six4-Moe::GFP, and five points were chosen along the interface confronting a dividing GSC (visualized by His2Av::mRFP1 and identified by condensed chromosomes), with points taken from two or more z-steps. The extracted x, y, z values were used as input in orthogonal distance regression to derive a best-fit plane. Second, for dividing GSCs, we extracted two sets of x, y, z coordinates to represent the division trajectory for anaphase, telophase, or metaphase nuclei. The coordinates were taken by marking the vertex of each chromosome complement, or for a metaphase figure, by marking a point on either side of the chromosome complex located midway on its long axis. The division angle was calculated as the angle between the best-fit plane and this inferred division trajectory. In all cases, division angle distribution between controls and ROKi-treated gonads were compared via a KS-test.

We also tested if control or ROKi divison angle distributions deviated from what might be expected as random. Note that from the perspective of an external reference plane, such as the niche-GSC interface, for a spherical cell, not all division angles are equally likely in a random model. If spindle position is not regulated in the cell, the distribution of spindles will exhibit a bias toward shallow angles, as shown in Figure S2E, F, and as described in ref^38^. This is because the probability of orientation falling within a particular range is dependent on the fraction of the inside cortical surface available for spindle attachment. There is much more surface area to attach to away from the reference interface. Hence, more representation of lower division angles. We therefore calculated the probability of cells dividing within 10-degree intervals from 0-90 degrees, following the equation from ref^38^. We then graphed the actual percentage of control or ROKi cells that divided within these intervals compared to the expected percentages of a random distribution.

## Supplemental Information

Document S1. Figures S1-S4

