## Supplemental Information for "An actomyosin network organizes niche morphology and responds to feedback from recruited stem cells"

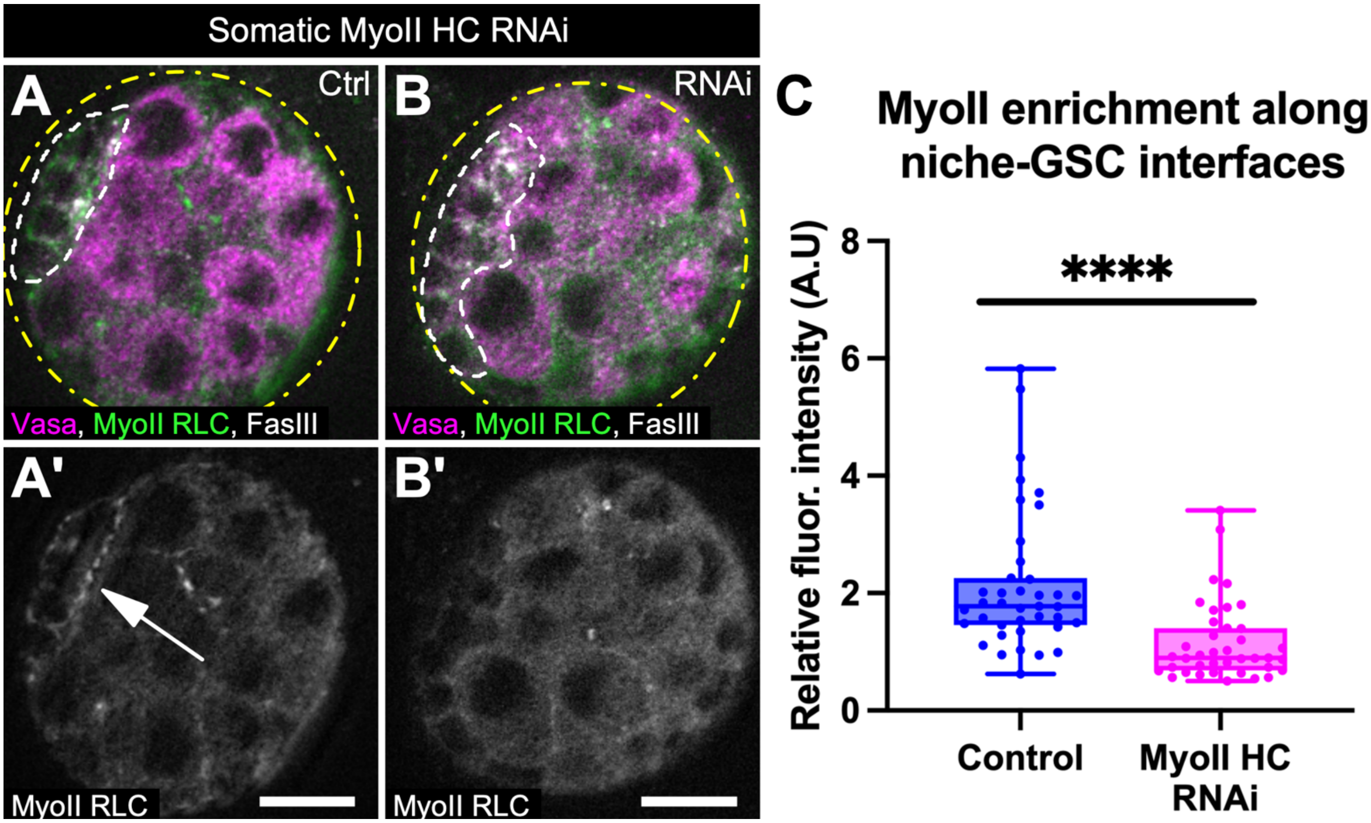


**Figure S1: MyoII is successfully depleted upon MyoII HC RNAi expression. Related to Figures 4 and 5.**

A-B) Control and MyoII HC RNAi gonads expressing a transgenic MyoII RLC::GFP construct stained for Vasa (magenta), GFP (green), and FasIII (white). A'-B') Single channel of MyoII RLC::GFP shows enrichment in controls towards niche-GSC interfaces (A', arrow), and reveals depletion in the niche of MyoII HC RNAi gonads (B'). C) Quantifications show MyoII HC RNAi in the somatic gonadal cells leads to decreased MyoII at the niche-GSC interface (****$p<0.0001$, Mann-Whitney).

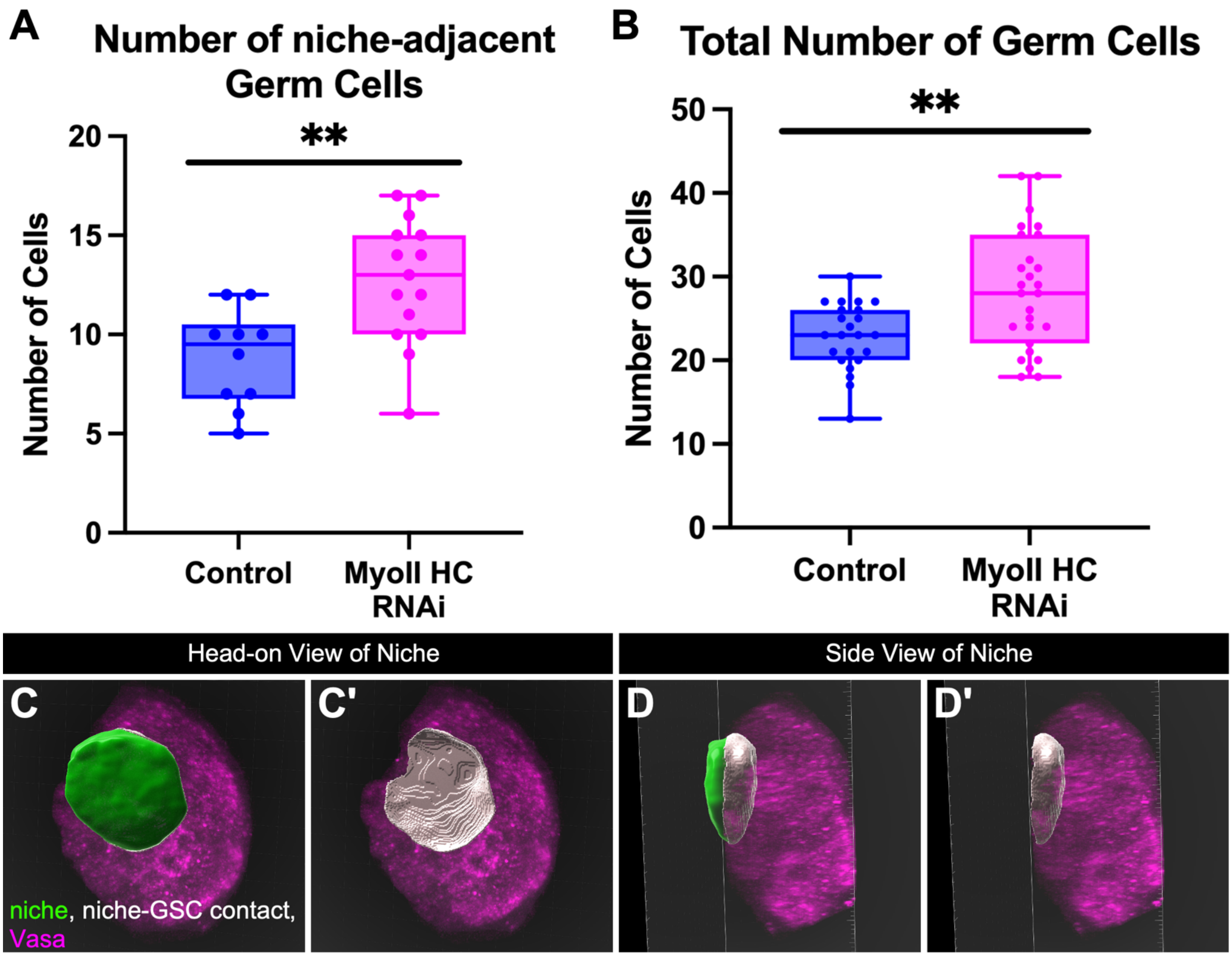


**Figure S2: Proper niche shape is required to limit contact with neighboring GSCs. Related to Figure 5.**

A) Number of germ cells, regardless of STAT activity, that contact the niche is increased in Six4-GAL4 > UAS MyoII HC RNAi gonads compared to sibling controls. B) Total number of germ cells in the Six4-GAL4 > UAS-MyoII HC RNAi gonads is increased compared to sibling controls. (For A and B, **$p<0.01$, Mann-Whitney). C-D) Snapshots of a control gonad 3-D projection from Imaris with Vasa visualized in magenta. The green surface represents the niche, and the grey surface represents the surface area of the niche that contacts the germline. C-C') The gonad is oriented so the niche surface is viewed head-on. D-D') The gonad from panel C is rotated ~90 degrees left to view the niche from the side. C'-D') The niche surface is turned off so the niche-GSC contact area can be viewed alone relative to the germline.

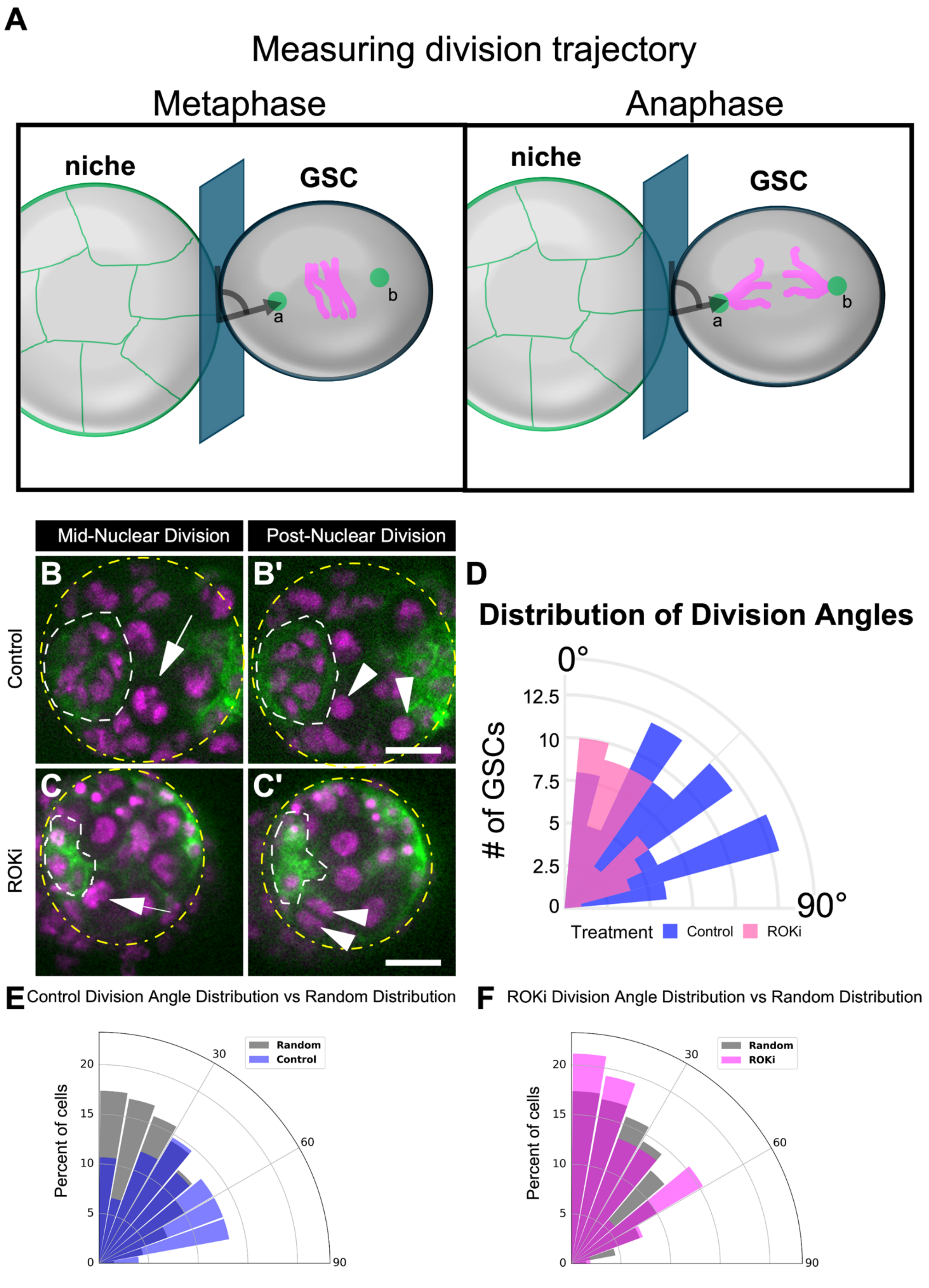


**Figure S3: Proper niche shape ensures non-random GSC division orientation. Related to Figure 5.**

A) Schematic of division angle analysis using His2AV::mRFP1 to visualize condensed chromosomes (pink). A plane of best fit (blue) was generated between the niche and a given GSC along their contact surface. The left panel shows how we established division trajectory for a metaphase GSC by extracting two points, a and b, on either side of, and perpendicular to, the metaphase plate. We calculated the angle line ab makes relative to the plane of best fit. The right panel shows how we extract division trajectory for an anaphase GSC by marking two points, a and b, on the separating chromosomes. Division trajectory was calculated by measuring the angle that line ab makes with the plane of best fit. In both examples, we show a GSC dividing generally away from the niche (with an angle close to 90 degrees). B-C) Live-images of Six4-Moe::GFP, His 2::RFP gonads to track nuclear division orientation of control (B) or ROKi treated (C) cells. B) Early anaphase nucleus (arrow). C) Prometaphase nucleus (arrow) B’-C’) Late telophase nuclei of the respective cells from panels Band C (arrowheads). D) Distribution of division angles, binned in 10-degree increments, from untreated (blue symbols, n=75 nuclear divisions) or ROKi treated gonads (magenta symbols, n=52 nuclear divisions; ****p<0.001, KS Test). E-F) Graphs of the percentage of total control (E, blue wedges) or ROKi (F, magenta wedges) cells dividing within 10 degree intervals from 0-90 degrees. Grey wedges represent the expected random division angle distribution of spherical cells.

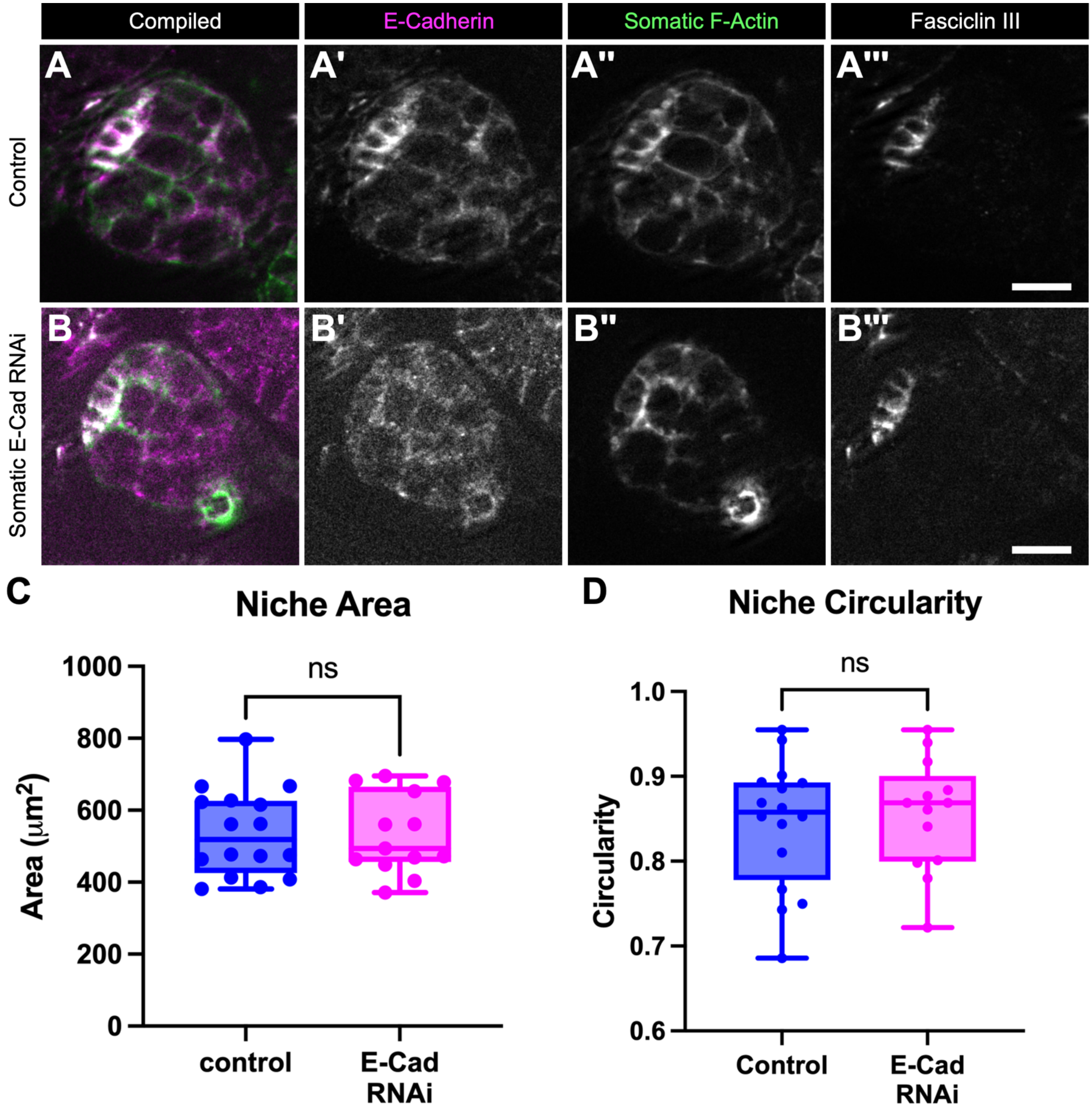


**Figure S4: E-Cadherin is not required for niche compaction. Related to Figure 7.**

A-B) Six4-Moe::GFP Control (A) and Six4-GAL4 > UAS-E-Cad RNAi gonad with E-Cadherin (magenta, A'-B'), Somatic F-actin (Green, A''-B''), and Fasciclin III (White, A'''-B''') labeled. Note the localization of E-Cad in the control niche (A'), but the depletion of E-Cad in the RNAi gonad (B'). C-D) Niche area (C) and niche circularity (D) are unchanged in E-Cad RNAi gonads compared to sibling controls (Mann-Whitney).
