## Supplementary material for "An actomyosin network organizes niche morphology and responds to feedback from recruited stem cells": Reagents Table

### Key resources table

| REAGENT or RESOURCE | SOURCE | IDENTIFIER |
|---|---|---|
| Antibodies | | |
| Goat polyclonal anti-Vasa | Santa Cruz Biotechnology | Cat# sc-26877 (dC-13), RRID:AB_793880 Discontinued |
| Mouse monoclonal anti Fasciclin III | Developmental Studies Hybridoma Bank | DSHB:7G10; RRID:AB_528238 |
| Rabbit polyclonal anti STAT92E | E. Bach | N/A |
| Rabbit polyclonal anti RFP | Abcam | ab62341; RRID:AB_945213 |
| Mouse monoclonal anti Gamma Tubulin | Sigma | GTU-88, T6557, RRID:AB_477584 |
| Chick polyclonal anti GFP | Aves Labs | Cat#GFP-1020; RRID: AB_2307313 |
| Rat monoclonal anti DE-Cadherin | Developmental Studies Hybridoma Bank | DSHB:DCAD2 RRID: AB_528120 |
| Normal Donkey Serum (NDS) | Jackson ImmunoResearch | Cat#: 017-000-121; RRID: AB_2337258 |
| Alexafluor Secondary Antibodies (488, 647) | Molecular Probes | |
| Cy3 Affinipure Secondary Antibodies | Jackson ImmunoResearch | Cat#: 711-165-152; RRID: AB_2307443 |
| Chemicals, peptides, and recombinant proteins | | |
| Para-Formaldehyde (PFA), 16% | Electron Microscopy Sciences | Cat#15710 |
| Hoechst | Sigma | Cat#33342; CAS Number: 875756-97-1 |
| Propyl-gallate | Sigma Aldrich | SKU: P3130; CAS Number 121-79-9; PubChem Substance ID 24898394 |
| Normal Donkey Serum (NDS) | Jackson Immunoresearch Laboratories | 017-000-121; RRID: AB_2337258 |
| Ringer's solution | Other | doi:10.1101/pdb.rec12409 |
| Schneider's Insect Media | GIBCO | 21720-024 |
| Insulin, bovine | Sigma | Cat# 10516 |
| Penicillin/ Streptomycin | Corning | 30-002-CI |
| Fetal Bovine Serum | GIBCO | Cat# 10082 |
| Critical commercial assays | | |

| | | |
|---|---|---|
| Gateway LR Clonase II Enzyme mix | ThermoFisher Scientific | Cat#: 11791020 |
| SpinSmart Plasmid Purification: Low-copy plasmid DNA, P1 constructs, or cosmids | Denville Scientific | Cat#CM-410-50 |
| **Experimental models: Cell lines** | | |
| DH5 alpha Cells | NEB | Cat#C2987H |
| oneShot Top10 | Invitrogen | Cat#C404006 |
| **Experimental models: Organisms/strains** | | |
| P-Dsix4-eGFP::Moesin | R. Lehmann | Sano et la 2012 |
| nos-moesin::GFP | R. Lehmann | Sano et al 2005; FBtp0040584 |
| UAS-Stat92E RNAi | Bloomington Drosophila Stock Center | Flybase: FBst0033637; RRID:BDSC_33637 |
| sqh-Sqh::mCherry | A. Martin | Rauzi et al., 2010 |
| sqh-Sqh::GFP3x | Y. Bellaïche | Pinheiro et al., 2017 |
| UAS-GFP::Zipper | D. Kiehart | Curr. Biol. 15, 2208–2221 |
| UAS-F-Tractin::tdTomato | w[*]; P{w[+mC]=UASp-F-Tractin.tdTomato}15A/SM6b; MKRS/TM2 | RRID:BDSC_58989 |
| Six4-Gal4 | S. DiNardo | Anllo et al., 2019 |
| Six4-Gal4::VP16 | S. DiNardo | This Work |
| nos-Gal4-VP16 | R.Lehmann and E. Selva | Van Doren et al., 1998 |
| P{CaryP}attP2 | Bloomington Drosophila Stock Center | FBti0040535; RRID:BDSC_8622 |
| His2Av::mRFP1 | Bloomington Drosophila Stock Center | FBtp0056035 |
| UAS-MyoII HC RNAi | Bloomington Drosophia Stock Center | RRID:BDSC_65947 |
| UAS-MyoII RLC RNAi | Bloomington Drosophia Stock Center | RRID:BDSC_33892 |

| | | |
|---|---|---|
| UAS-Cdc25 RNAi | Bloomington Drosophia Stock Center | RRID:BDSC_34831 |
| P(UAS-hid.Z)2 | Bloomington Drosophia Stock Center | RRID:BDSC_65403 |
| P{w[BmC]=Dfd-EYFP.w[BmC]}2 | Bloomington Drosophia Stock Center | RRID:BDSC_8623 |
| TM6B, P{Dfd-GMR-nvYFP}4, Sb[1] Tb[1] ca[1] | Bloomington Drosophia Stock Center | RRID:BDSC_23232 |
| UAS-E-Cadherin RNAi (Shg RNAi) | Bloomington Drosophila Stock Center | RRID:BDSC_32904 |
| **Oligonucleotides** | | |
| Lower strand, Forward CACCCAGCAAAGACCGTGAGTTG | Adapted from Clark et al, 2006 | |
| Upper strand, Reverse GTTGGATCCATTGCCATCCAGTTG | Adapted from Clark et al, 2006 | |
| **Recombinant DNA** | | |
| pENTR/D-Topo entry vector | Invitrogen | K240020 |
| pBPGAL4.2::VP16Uw | Addgene | Pfeiffer et al., 2010; RRID:Addgene_26228 |
| **Software and algorithms** | | |
| Imaris | https://imaris.oxinst.com/products/imaris-for-core-facilities | |
| Surface-Surface Area Contact Xtension | https://imaris.oxinst.com/open/view/surface-surface-contact-area | Matthew Gastinger |
| FIJI (Image J) | www.fiji.sc | RRID:SCR_003070 |
| Image J | www.imagej.nih.gov/ij/ | RRID:SCR_003070 |
| Metamorph Microscopy Automation and Image Analysis Software | Leica; https://www.moleculardevices.com/products/cellular-imaging-systems/acquisition-and-analysis-software/metamorph-microscopy | v7.8.40; RRID:SCR_002368 |
| Axio-Vision Imaging Software | Zeiss | v4.8.1 |

| | | |
|---|---|---|
| EZFig | Image J Plugin | Benoit Aigouy |
| Graphpad Prism | Graphpad Software | v7.0; RRID:SCR_002798 |
| BioRender | BioRender.com | |
| Other | | |
| Matek imaging dish | Thermofisher | P35G-1.5-14-C |
| Leica DM16000 B inverted spinning disk confocal | Leica | |
| 63x / 1.2 NA water immersion objective | Leica | 506279 |
| 40x / 1.1 NA water immersion objective | Leica | |
| Leica M165FC | Leica | |
| Leica M165C | Leica | |
| GFP Filter set ET470/ 40x; ET525/50m | Leica 10447408 | |
| mCherry Filter set ET560/ 40x; ET630/75m | Leica 10450195 | |
| DAPIFilter set AT350/ 50x; ET460/50m | Leica 10450196 | |
| Achromat 1.6x objective | Leica 10450163 | |
| Video 0.63x objective | Leica 10447367 | |
| AxioCam HRm | Zeiss | |
| 40x / 1.2 NA water immersion objective | Zeiss | |
| 20x / 0.8 NA objective | Zeiss | |
| Alfa Aesar Tungsten wire | Fisher Scientific | AA10408G6 |
| Needle holder | Fisher Scientific | 08-955 |
| Nytex basket | | |
